# Evolving copy number gains promote tumor expansion and bolster mutational diversification

**DOI:** 10.1101/2022.06.14.495959

**Authors:** Zicheng Wang, Yunong Xia, Lauren Mills, Athanasios N. Nikolakopoulos, Nicole Maeser, Jason M. Sheltzer, Ruping Sun

## Abstract

Charting the evolutionary history of rampant somatic copy number alterations (SCNA) is an indispensable step toward a deeper understanding of their roles in tumor development. However, the existing SCNA timing analysis is limited to low copy number states and initiating gains, which are not necessarily close to the onset of the malignant proliferation. Moreover, it remains a critical question if the timing of an SCNA reveals the corresponding variant’s fitness effect. Here we propose a framework to estimate the arrival time of a clonal SCNA, i.e., the time delay from its last alteration to the start of population expansion, in addition to its initiation time when the first alteration occurs. Our method leverages the bias that a genomic segment, when resting on a copy number (CN) state, accumulates somatic single nucleotide variants (SSNV) at a rate proportional to its CN. From the whole genome sequencing data of 184 samples from 75 patients across five tumor types, we commonly observed late clonal CN gains following early initiating events, which appear right before the expansion leading to the observed tumor sample(s). Genome doubling (GD) can be late, but post-GD CN evolution is prevalent in the genealogy of the most recent common ancestor of patient tumors. Notably, mathematical modeling suggests that late evolving events could contain rate-limiting drivers. The advantage of evolving gains could arise from the dosage increase of cancer genes in proliferative signaling and amplification of early functional variants. In addition, evolving SCNAs bolster the diversification of SSNVs between sub-populations, exacerbating the vicious circle between malignant growth and accumulation of genomic heterogeneity. Our findings have broad implications for resolving the trajectory of SCNAs, discerning the CN markers of malignant growth *in vivo*, and properly quantifying tumor heterogeneity in aneuploid tumors.

## 1 Introduction

Underlying the maintained genomic diversity within a patient tumor is the uncontrolled proliferation, a core hallmark of cancer [1], coupled with somatic alterations occurring over time [2]. To prevent the disease, uncovering the somatic aberrations responsible for the malignant growth is the primary goal of precision oncology. At the genomic level, somatic alterations exist on a spectrum, ranging from small changes such as somatic single nucleotide variants (SSNV, often referred to as mutations) [3] to large somatic copy number alterations (SCNA). Frequent chromosomal mis-segregation (chromosomal instability or CIN) leads to abnormal chromosome numbers (aneuploidy[4]) and unbalanced structural variations (SV) cause segmental SCNAs [5]. These two genomic errors are intertwined in many solid tumors, leading to extensive SCNAs, especially in advanced diseases [4] with poor clinical outcomes [6].

The inextricable relation of SCNAs to cancer initiation[7, 8] and progression[9] has become a consensus in cancer genomics. It remains little known, however, to what extent a specific SCNA accounts for the malignant growth and how it affects the intra-tumor-heterogeneity (ITH) [4]. Indeed, chaotic karyotype and widespread high copy number (CN) states in aneuploid tumors[10] make it extremely challenging to identify SCNA drivers, limiting the precision of using SCNA patterns for diagnostic and treatment purposes. For example, the treatment strategy for osteosarcoma, the most common bone tumor affecting teenagers with one of the most chaotic aneuploid genomes, has stagnated for decades [11]. From an evolutionary perspective, discovering the tempo of SCNA during somatic evolution is key to gaining knowledge of SCNA drivers [12]. Here, we hypothesize that the timing of SCNAs can be systematically measured from whole genome sequencing (WGS) data of patient tumors, and the temporal axis contains tangible information in isolating the effect of specific SCNAs on tumorigenesis.

We should pause to clarify how bulk sequencing data capture somatic evolution timeline. The tumor founder cell arose from the succession of clonal expansions in the pre-cancerous context where beneficial alterations endow progenitor cells with the ability to crowd out less advantageous populations [13] (Figure 1A). The growth of the primary lesion gives rise to genomically diverging lineages [14], some of which, after acquiring a more malignant potential, can initiate the re-growth of a secondary tumor, such as metastasis [15]. Bulk sequencing data provide us with the opportunity to anchor the roots of expansion (the most recent common ancestor, or MRCA). Clonal variants in a single sample refer to the root of the observed sample. In multi-region sequencing, truncal variants from multiple samples could trace back asymptotically to the founder of the tumor [16]. For longitudinal sampling, e.g., of paired primary and metastatic tumors, truncal variants could point to the MRCA of the branched tumor progression [17]. Multi-samples reflect the population expansion at a broader scale, i.e., they coalesce to an earlier progenitor cell than a single sample does. Collectively, truncal variants revealed by a particular sampling strategy must map to the somatic evolution timeline prior to the corresponding *sampling-relevant expansion* (SRE).

**Fig. 1.**
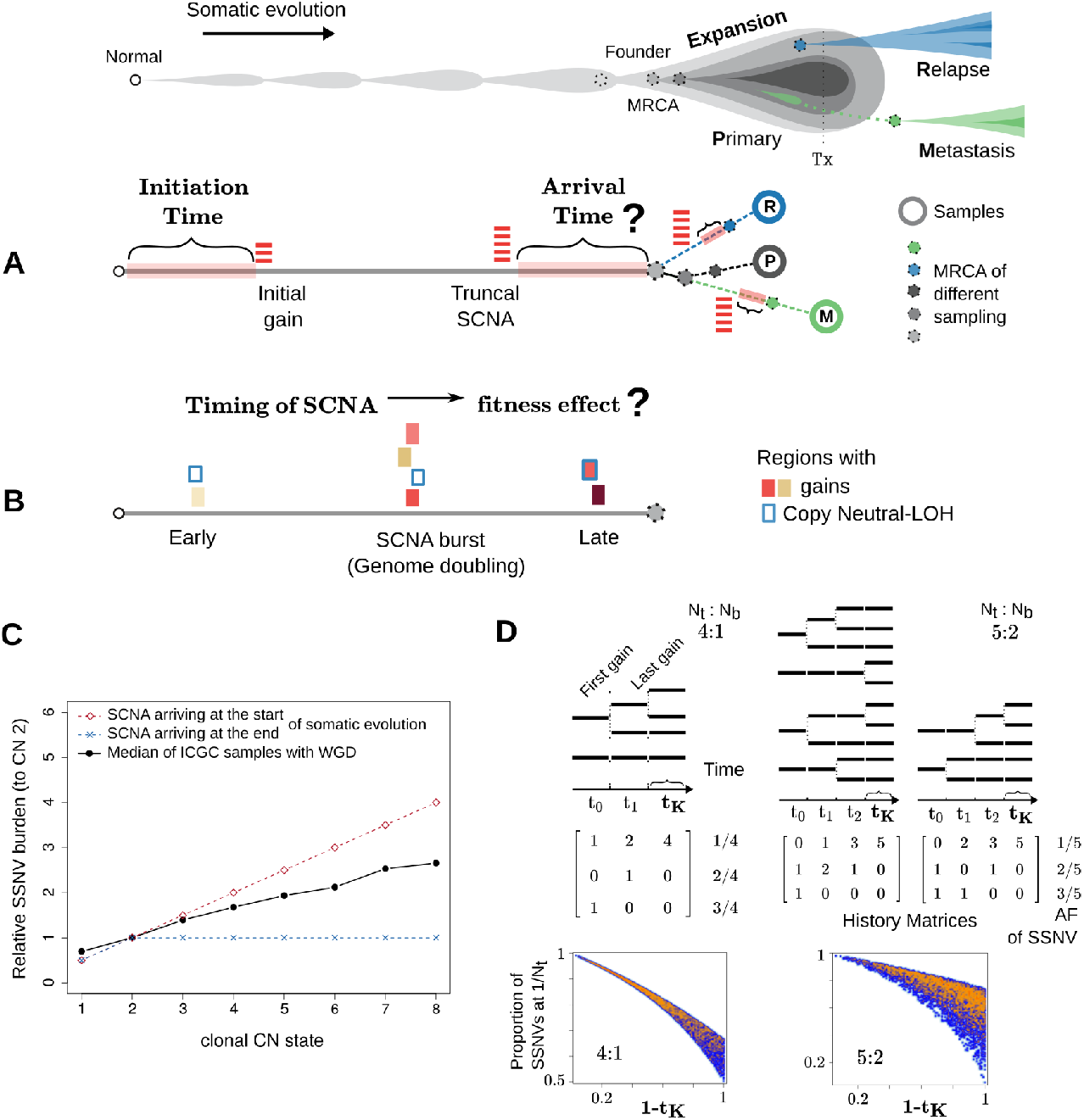
Measuring the arrival and initiation time of SCNAs. A drawing at the top marks the concept of cancer somatic evolution, which leads to the establishment of the most recent common ancestor (MRCA) of the primary tumor, as well as the roots of secondary tumors. Tx: treatment. (A) For an SCNA present within the MRCA of a sampling-relevant tumor expansion, we aim to characterize the time when the last gain and first gain appeared (referred to as arrival and initiation time, respectively) for the corresponding genomic region. (B) We further aim to address the question if the late truncal gains are neutral or beneficial to tumor evolution. (C) The solid black line shows that the burden of SSNVs of a given genomic locus correlates with its CN state in tumors sequenced by ICGC (single sampling, with WGD). Two dashed lines assume the two extreme scenarios of SCNA arrival time. (D) The proportion of SSNVs at different allele fractions depends on the SCNA history matrix and the relative time span of each CN stage. The smoothed scatter plots show the burden of single-copy SSNVs against the arrival time of an SCNA (*t*_*K*_), with an increasing density from blue to orange. Two example SCNA states are shown (with CN configuration at 4:1 and 5:2, respectively).

The timing of a truncal SCNA (more specifically, gains) on the evolution toward the MRCA could shed light on the impact of this variant on promoting the SRE. However, our knowledge about the SCNA timing remains fragmentary as the existing methods are restricted to simple (single or double) gains [16, 18, 19]. Single-sample based pan-cancer analysis from ICGC[19] revealed extensive variation in the molecular time of these sample gains. Relative to aneuploid tumors, these gains are likely the initiating events of copy number evolution and may not be sufficient to induce the expansion. For example, single chromosome gains have shown limited capability in driving proliferation in *in vitro* systems [20]. As such, it would be critical to know when a genomic segment further evolves beyond the initial gains, which can often culminate at a state greater than four copies [10]. This requires a timing method applicable to high SCNA states. In SCNA timing analysis, the following assumption is made: the site frequency spectrum (SFS) of SSNVs in a genomic region affected by SCNA depends on the trajectory (the order) and time span on each CN state that the segment had ever rested on [21]. For a single gain, the ratio between early (duplicated) and late (non-duplicated) SSNVs can be used to estimate the relative timing of the gain [12]. For a high CN state (*>* 4) or gains involving both maternal and paternal alleles, the detailed trajectory of each SCNA evolution is not identifiable from the SFS [18], at least not without further constraints on the trajectory or additional information that could allow one to reconstruct the ordering [21]. We reason, however, that it is possible to obtain the upper bounds of the time duration of a segment resting on its observed truncal CN state before the onset of SRE. In principle, once a segment arrives at the final CN state, it can only accumulate single-copy SSNVs; the longer the segment persists in the observed CN state, the more overwhelming the single-copy SSNVs. Notably, for the first time, this allows us to investigate the *arrival time* for SCNAs at high CN states.

An outstanding question is how the timing of an SCNA reflects its fitness effect (Figure 1B). Whereas early gains could initiate and increase the risk of disease, we argue that late-appearing SCNAs close to MRCA could promote the population expansion more directly. If an SCNA triggers SRE, one may foresee that the progenitor cell proliferates vigorously upon acquiring the corresponding SCNA. By contrast, before the SRE begins, if the clonal lineage persists for many generations after receiving an SCNA, the corresponding variant is unlikely to be sufficient to drive the SRE. Punctuated acquisition of polyploidy (e.g., through genome doubling or GD) is prevalent in aneuploid tumors [22] but it remains unclear how close the occurrence of GD is to tumor transformation. Evidence exists that GD itself doesn’t confer a strong fitness advantage[23]; instead, it can enhance the plasticity of the genome that permits further CN evolution, such that aneuploid cells adapt to overcome possible fitness penalties incurred by GD [24]. Therefore, SCNAs that arrive after GD could contain driver events. Moreover, depending on the precise location of biopsied tissue, single-sample analyses may differ in the corresponding time scale; subsequently, it is particularly essential to focus on the timeline toward the malignant growth - the somatic evolution in collecting truncal SSNVs of multiple samples of a tumor (e.g., multi-region samples or paired primary and metastatic samples).

To broaden the “timable” genomic regions for SCNAs, we developed `Butte` (BoUnds of Time Till Expansion), a computational framework to estimate the upper bounds of SCNA arrival and initiation time from WGS data. By applying `Butte` onto multi-sampled WGS data of five cancer types with widespread SCNAs, including osteosarcoma, we systematically charted the temporal patterns of CN evolution *in vivo*. To see if late-appearing SCNAs may confer fitness benefits, we constructed mathematical models to examine the evolutionary mechanisms that give rise to these late truncal events. Furthermore, we also interrogated potential ways the late culminating SCNAs could add to the fitness and revealed its impact on mutational diversification during tumor expansion.

## 2 Results

### 2.1 A computational framework to estimate the arrival time of SCNA gains

From the WGS data, one can characterize with high certainty the dominant SCNAs, inferring the integer allelic CN of a genomic region and the cellular prevalence of the corresponding SCNA, i.e., the percentage of cells sharing the dominant SCNA state [25]. We refer to a unique version of a genomic region (or segment) as an “allele”. We term the total CN as *N*_*t*_ and the CN for the minor allele as *N*_*b*_ (“b” stands for b-allele determined by germline SNPs) for a dominant SCNA. The “allele” fraction of an SSNV is the amount out of the total *N*_*t*_ copies of the region that carry the corresponding variant. We found that in the aneuploid tumors sequenced by ICGC (International Cancer Genome Consortium), the SSNV burden increases with the dominant SCNA states of the corresponding genomic region (Figure 1C). Such a pattern is due to an intrinsic bias between SCNAs and SSNVs: a genomic segment resting on a CN state accumulates SSNVs at a rate proportional to the corresponding CN. Thereby, the burden and CN multiplicity of SSNVs are actively shaped by SCNAs.

Generally, the observed SCNA of a genomic segment (with a configuration *N*_*t*_ : *N*_*b*_ different from 2 : 1) is the result of a series of CN events. For an SCNA involving at least *K* gain events, the total time of somatic evolution can be divided into *K* + 1 stages. The genomic segment begins with the 2 : 1 setting in the first stage and keeps “climbing up” by duplicating one of its existing copies in each subsequent stage, respectively, until it arrives at the observed SCNA state in the last stage (Figure 1D). Accordingly, each stage is associated with certain time proportion (*t*_*k*_ ≥ 0) and 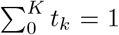, the total time for the somatic evolution. SSNVs occurring at stage *k* on a segment copy that experiences duplication(s) in later stages will be presented on the duplicated copies (*a* ≥ 2*/N*_*t*_). By contrast, SSNVs acquired at stage *k* on a copy without further duplication events remain at the single allele state (*a* = 1*/N*_*t*_). One can define a history matrix *A* with entry *A*_*jk*_ representing the number of segment copies in stage *k* that result in a final allele state *a*_*j*_ [18, 21]. It can be seen that the abundance of SSNVs at allele state *a*_*j*_ depends on Σ *A*_*jk*_*t*_*k*_. From the site frequency spectrum (SFS) of SSNVs in a region affected by SCNA, one can estimate the relative abundance of SSNVs at each allele state, and in turn, solve for each *t*_*k*_. There has been much effort to infer *t*_0_, i.e., the timing of the first copy number event [18, 19]. These efforts focused on single gain (2 : 0 and 3 : 1) and at most double gains (3 : 0 and 4 : 1), where the history matrix *A* is invertible. By contrast, for other SCNA states, multiple possible trajectories can exist and the underlying linear system is underdetermined, i.e., there are more time stages (unknown variables) than the possible allele states (equations). We note that, however, regardless of the underlying history, multi-allele SSNVs (≥ 2*/N*_*t*_) can only occur before the last stage (*K*) of CN evolution; once the genomic region arrives at the observed clonal SCNA state, all the copies (*N*_*t*_) would accumulate SSNVs at single allele state (1*/N*_*t*_). Therefore, the longer the last stage of CN evolution (from the emergence of the clonal SCNA to the onset of population expansion), the more overwhelmingly the single allele SSNVs dominate the SFS (Figure 1D). Such monotonicity allows one to calculate the bounds of the time duration for the last stage even when the detailed SCNA history is unidentifiable.

To investigate how various SCNAs unfold during somatic evolution, we developed `Butte` (<u>B</u>o<u>U</u>nds of <u>T</u>ime Till <u>E</u>xpansion), which adopts linear programming to infer the upper bounds of arrival (*t*_*K*_) and initiation time (*t*_0_) of SCNAs (see Methods). From a technical point of view `Butte` extends the full maximum-likelihood estimation procedure implemented in `cancerTiming` [18]. Notably, `Butte` does not restrict the analysis to single and double gains, but in addition allows the calculation of the upper bounds of *t*_*K*_ and *t*_0_ for SCNAs up to seven total copies, broadening the “timable” SCNA regions. The upper bounds, by definition, overestimate the time duration. Nevertheless, these estimates systematically correlate with the actual timing of SCNA initiation and culmination (Supplementary Figure 1). To test the performance of `Butte` on real patient tumors, we first evaluated the timing predictions by analyzing multi-region WGS data of colorectal adenocarcinomas (COAD) [16, 26]. `Butte` successfully identified early CN gain of chromosome (chr) 5q (Supplementary Figure 2), corresponding to the SCNA state of 2 : 0 (copy neutral loss of heterozygosity), a known early step in COAD initiation involving gene *APC* [27]. As a benchmark for late-appearing events, private (sample-specific) SCNAs should contain events that occur in the descendent lineage of the MRCA of multi-samples. `Butte` predicted their arrival time to be later than the public SCNAs on the timeline toward the MRCA, highlighting its capability in revealing the late arriving SCNA events (Supplementary Figure 3).

### 2.2 Evolving SCNA gains define the tumor transformation leading to the most recent clonal expansion

To evaluate the tempo of SCNAs in solid tumors, we applied `Butte` on five tumor types by analyzing eight published WGS datasets: osteosarcoma (OS) [28, 29], breast invasive carcinoma (BRCA) [30, 31], colorectal adenocarcinomas (COAD) [16, 26], esophageal carcinoma (ESCA) [32], and prostate adenocarcinoma (PRAD) [33], six of which comprise multi-sampling of patient tumors (Figure 2, Table 1). 70% of the analyzed genomes (corresponding to 87% of the patients) were near triploid, with the median fraction (IQR) of the high amplitude CN regions (≥ 4) being 0.37 (0.23 to 0.49). Loss of heterozygosity (LOH) is prevalent but mostly is at copy neutral or amplified states in the triploid tumors. High amplitude gains can be recurrent across cancer types (e.g., chr 8q) or within a specific tumor type (e.g., chr 1q for BRCA, chr 17p for OS, and chr 7 for COAD). These recurrent gains presumably contain driver events [34], yet their tempo in somatic evolution remains uncharted. Notably, karyotypes largely remain stable across different samples of the same tumor, despite the presence of continued subclonal CN diversification in a relatively minor fraction of the genome.

**Table 1.**
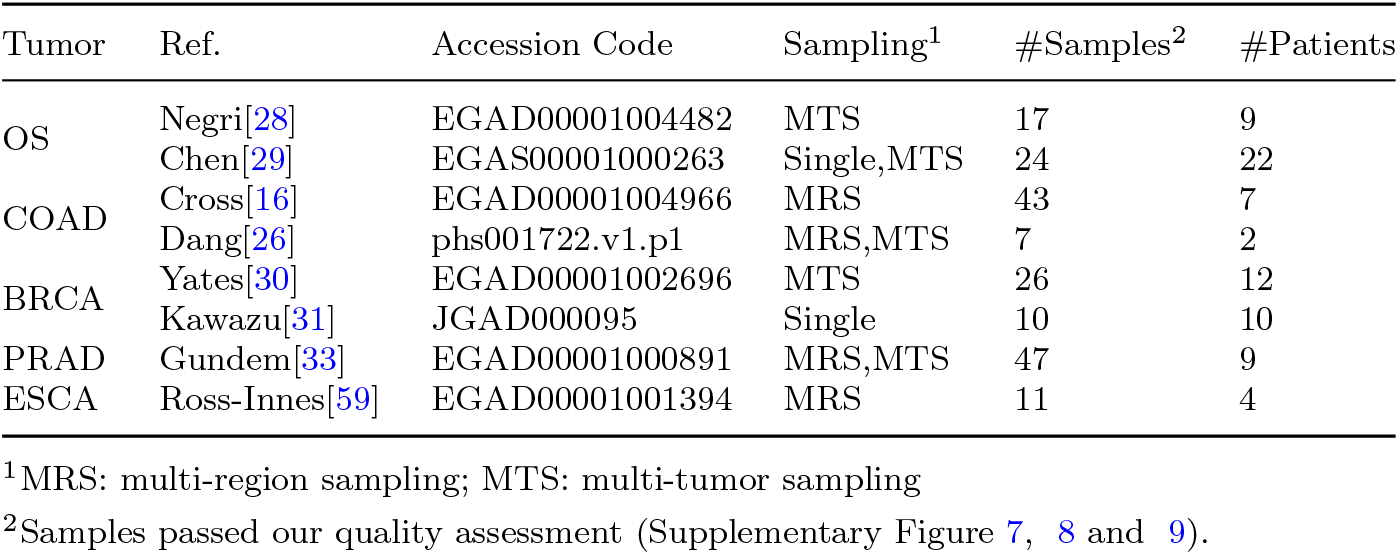
WGS data included in this study

**Fig. 2.**
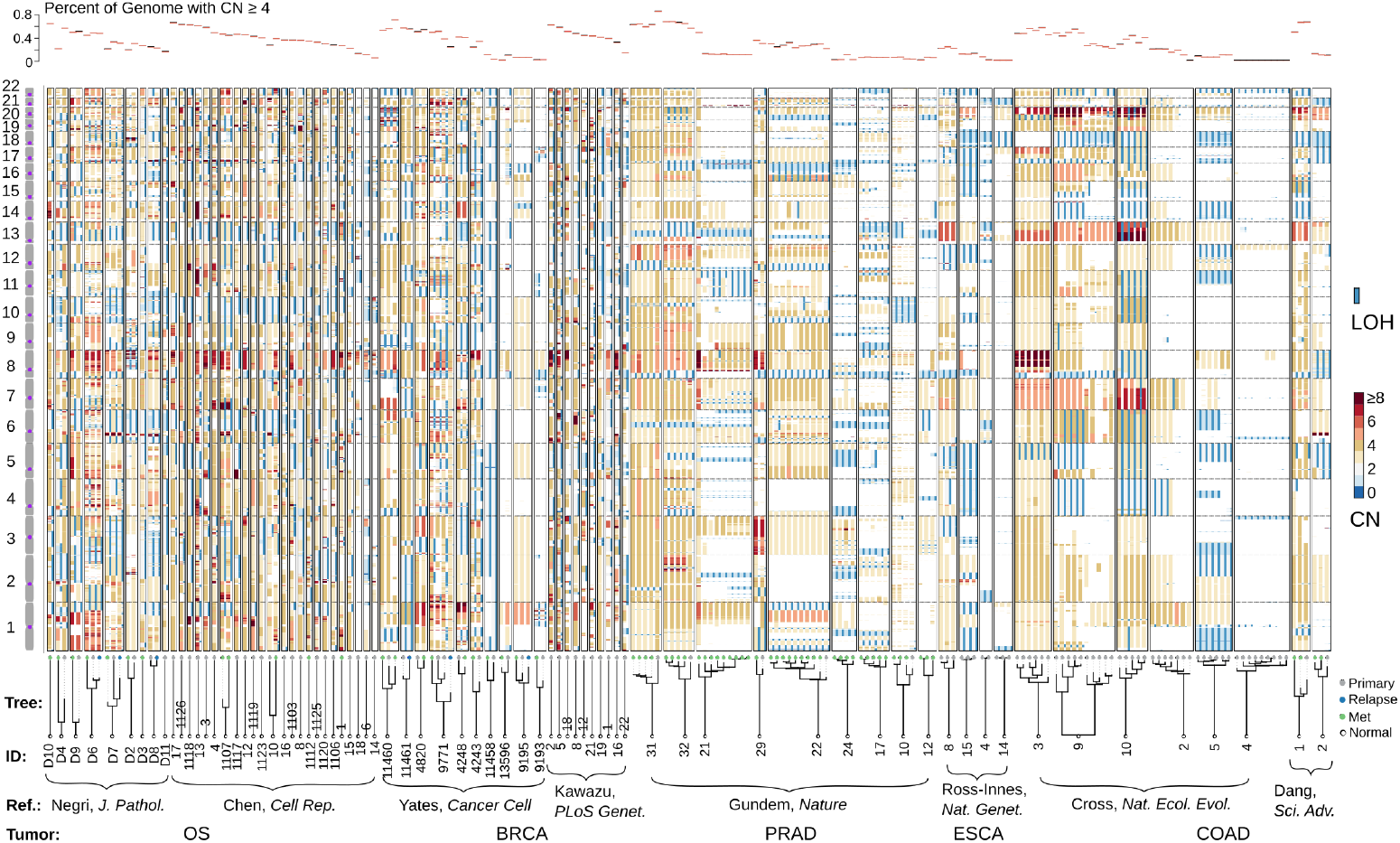
CN profiles across five tumor types from the re-analysis of published WGS data. Vertical bars represent the segmental CN states along the autosomal chromosomes characterized by the WGS data of tumor samples. For each sample, a color-coded thick bar shows the total CN state of each genomic locus and a thin blue bar to the right indicates that region has loss of heterozygosity (LOH). Samples belonging to the same patient are boxed. The top panel highlights the fraction of high CN states in each sample’s genome. The lower panel exhibits the sample phylogenetic trees constructed from SSNVs. Sample IDs, the reference where the WGS data was published, and tumor types are tabulated at the bottom. In this manuscript, a tumor sample is named after the concatenation of the tumor type, the first character of the author’s surname and the patient ID.

We note that 74 out 75 patient tumors acquired late-appearing gains close to MRCA regardless of the overall ploidy or tumor type (Figure 3A, B), with the only exception of COAD C 4, which shows high microsatellite instability. Punctuated copy number bursts were observed in the triploid samples, reflecting the ability of the genome to leapfrog over intermediate states to reach moderately high CN states through whole or partial genome doubling (GD)[35, 36]. Whereas GD occurs late (close to the MRCA) in some adult cancers (18 out of 34 patients), it appears to be an earlier event in many other tumors. This is particularly evident in OS where 28 of 30 patients had GD at the mid-stage of somatic evolution toward SRE (Figure 3B). The contrasting tempo of GD suggests that it probably has a context-dependent fitness. In tumors with early GD, `Butte` can characterize the post-GD CN evolution, whereby progenitor cells continue to sample the aneuploid fitness landscape [24]. Such an SCNA evolution involves stochastic chromosomal or structural abnormalities; however, certain genomic regions preferentially exhibit late gains across different patients in a particular tumor type, which, surprisingly includes those recurrent high amplitude gains, such as chr 8q in OS (Figure 3C) and chr 7 in COAD (Supplementary Figure 2). On the other hand, recurrent SCNAs appear to initiate early, e.g., chr 1q (*AKT3*) in BRCA, chr 8q and chr 17p (*TP53*) in OS and chr 5q (*APC*) in COAD (Figure 3C and Supplementary Figure 2). These additional gains pre- and/or post-GD could result from either high evolvability of the corresponding region, or persistent selection upon driver genes within.

**Fig. 3.**
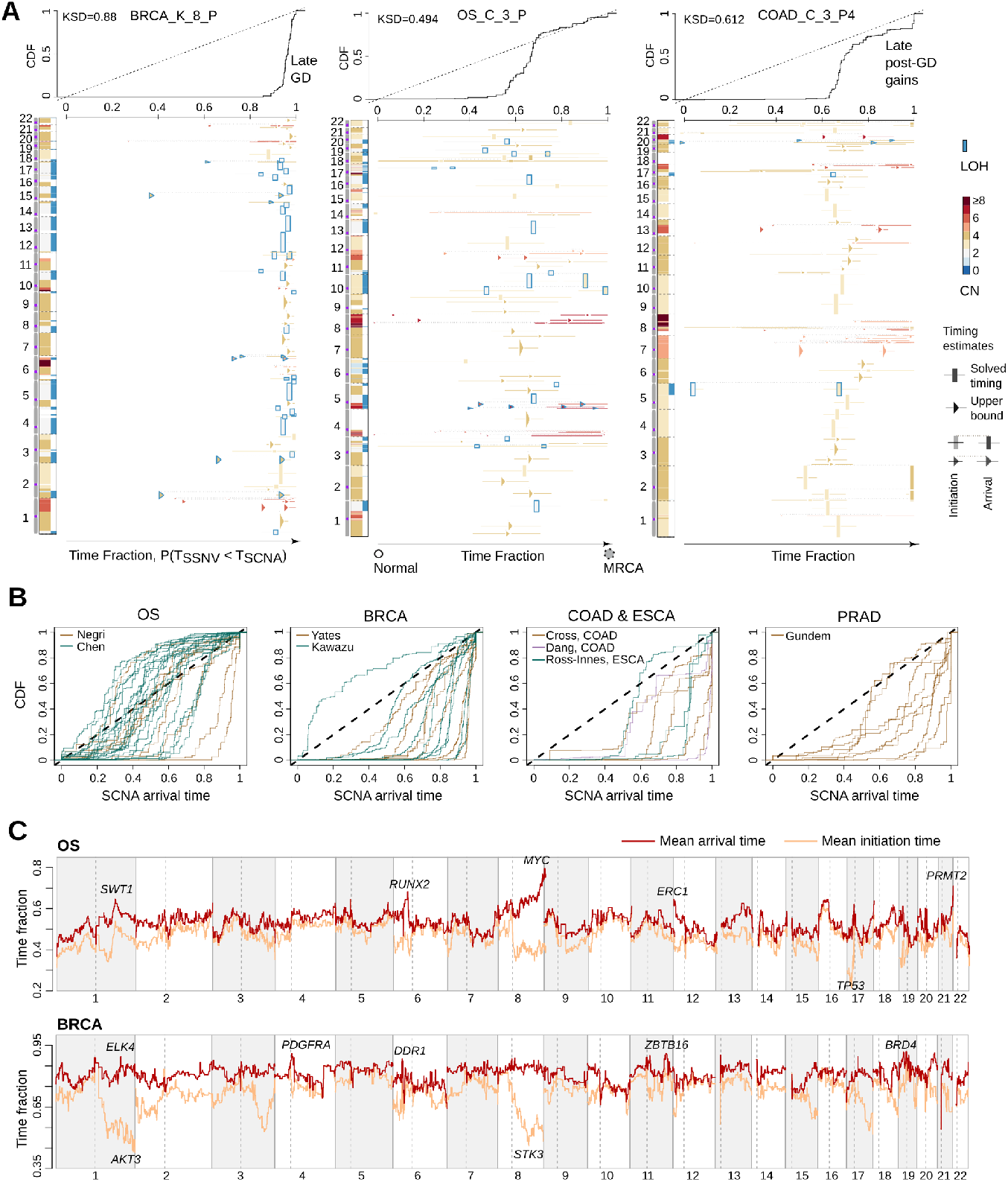
The timing patterns of SCNAs across five tumor types. (A) Timing plots of three exemplified tumors. CN states along the genome are shown on the left of each panel. The right panel visualizes the time fraction of somatic evolution from germline to the MRCA of the patient tumor. For each SCNA segment, the inferred timing is drawn by either rectangle (exactly solved timing) or arrows (upper bounds of timing when the solutions are not unique) with the same color-coding as its CN. The top panel shows the cumulative distribution (CDF) of SCNA arrival time. (B) The CDF curve of SCNA arrival time is shown for each patient categorised by the tumor type. (C) The average timing (both initiation and arrival time) for each one Megabase genomic bin across OS and BRCA patients are shown, respectively. Example genes in regions showing late arrival or early initiation times are marked.

The earlier the timing of GD, the more post-GD CN gains (Figure 4A). The late evolving gains are shorter in segment length than those associated with GD (Supplementary Figure 4), except for a few patients where post-GD events are sparse (e.g. for patient PRAD_G_21 and PRAD_G_31). This suggests that the post-GD CN evolution is driven by SVs, which occur at a higher rate than chromosomal mis-segregation. Indeed, the breakpoints of structural variants almost locate the boundary of SCNA segments (Supplementary Figure 5). As SVs continued to occur, it could become more focally amplified for the driver genes that are repetitively selected, making them more apparent in the late appearing gains, e.g., *MYC* [37] and *RUNX2* [38] in OS (Figure 3C). In terms of types of gains, amplified LOH (*N*_*b*_ = 0 and *N*_*t*_ *>*= 3) tend to culminate later than other types of amplifications, such as allele specific amplifications (*N*_*b*_ = 1 and *N*_*t*_ *>*= 3, *p <* 3*e* − 5), and this cannot be explained by the overestimation of `Butte` (Supplementary Figure 6 and 1). Whereas truncal LOH were supposedly acquired before GD [15] causing the complete loss of tumor-suppressor activity, the late appearing gains of the only remaining allele may indicate that these regions potentially acquire dosage-dependent gain-of-functions [39].

**Fig. 4.**
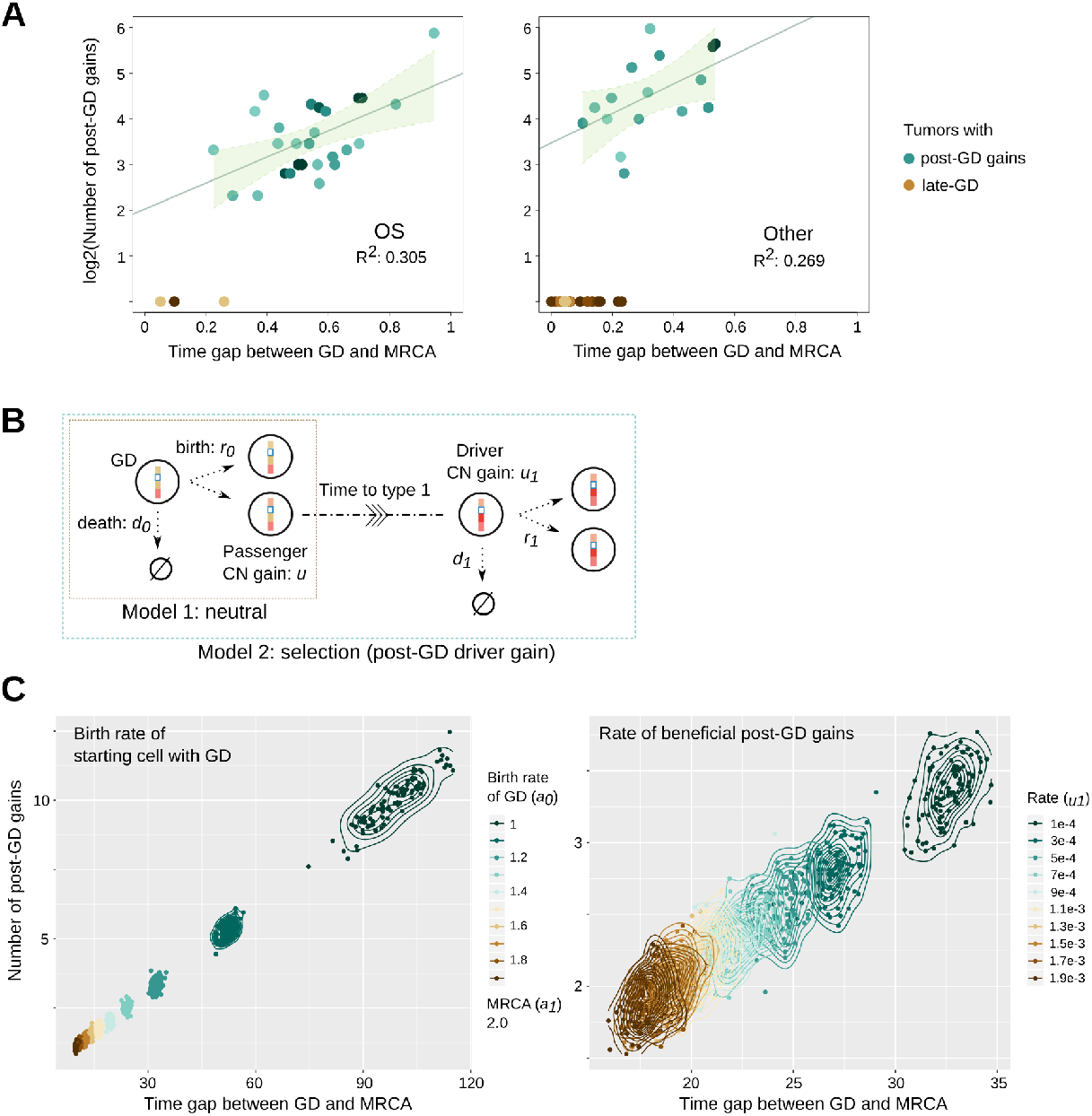
Mathematical modeling suggests that advantageous CN gains can occur late. (A) Scatter plots colored by density illustrate the number of post GD gains against the time fraction of post-GD evolution towards MRCA for OS and other adult cancer types, respectively. A linear regression line and the confidence intervals are indicated for tumors with post-GD gains. (B) The schema shows the setup of the two contrasting mathematical models: (1) GD is followed by neutral growth where additional gains do not confer a fitness advantage and (2) post-GD gains increase the growth rate. (C) Scatter plots display the number of post-GD passenger gains against the time of post-GD evolution characterized by the selection model. We studied the effects of the growth rate of GD (with a fixed growth rate of the MRCA, the left panel) and the rate of beneficial post-GD driver gains (the right panel), respectively. Each point refers to the average of 100 simulations. Contours of the 2d density estimation are shown. Parameters for the left panel: *b*_0_ = 1, *a*_1_ = 2, *b*_1_ = 1, *u*_0_ = 0.1, and *u*_1_ = 0.0001; right panel: *a*_0_ = 1.2, *b*_0_ = 1, *a*_1_ = 2, *b*_1_ = 1, and *u*_0_ = 0.1.

### 2.3 Mathematical modeling suggests the role of late gains in promoting tumor expansion

While early genomic changes during somatic evolution are suggested to be functional initiating events, there is little known about the effect of late-appearing alterations (e.g., close to the MRCA). To reveal the mechanism that gives rise to the tempo of GD and the number of post-GD events, we mathematically modeled the somatic evolution based on a multi-type branching process (see for example [40]), starting with a single tumor-initiating cell that just acquired GD (Figure 4B). This cell gives birth at a rate of *a*_0_ and dies at a rate of *b*_0_ (with net growth rate *λ*_0_ = *a*_0_ − *b*_0_ *>* 0). During progression, daughter cells acquire a passenger post-GD gain with a rate of *u*_0_, which does not change the net growth rate. With a lower rate *u*_1_, they can also acquire a driver post-GD gain, leading to increased fitness (*λ*_1_ = *a*_1_ − *b*_1_ *> λ*_0_, see Method for details). Our objective is to characterize the number of post-GD gains that reach fixation or become dominant in the observed tumor under two contrasting scenarios: one without and the other with driver post-GD gains.

We first assumed that all post-GD gains are passenger variants. Conditioned on the non-extinction of the population, we obtained that the number of post-GD gains accumulated before the MRCA that grows into detectable tumor follows a geometric distribution with parameter 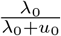 and mean 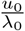. The mode of this distribution is at zero, similar to the cases where GD appears late and post-GD CN gains are rare. To tolerate the inclusion of subclonal but dominant SCNAs as the clonal variants, we further evaluated the dominant post-GD gains shared by the majority (≥ 90%) of cancer cells. Building on the results of [41], we derived the expected number of dominant post-GD gains in a tumor with size *N* as

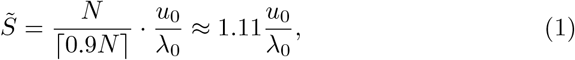

which is only slightly larger than the clonal ones. Assuming that *u*_0_ and *λ*_0_ are comparable, 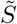 would be no more than just a few. Moreover, numerical simulations show that the number of dominant post-GD gains continues to follow a geometrical-like distribution with the mode at zero. Thus, if post-GD gains do not provide growth benefits, GD would be one of the last events before the MRCA as few of post-GD gains can become dominant in the observed tumor.

However, if post-GD gains increase fitness, the situation drastically changes. To emphasize how this happens, let us consider the first advantageous cell (type 1), which is introduced via a beneficial post-GD gain and grows into an infinite number of descendants. The original population without the new driver forms the type 0 population. In this scenario it suffices to focus on type 1 cells as their population most likely overshadows the type 0 population upon detection. The expected number of passenger post-GD gains 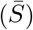 carried by a type 1 cell would be proportional to the time of occurrence of the type 1 cell. In Methods we show that the distribution of the birth time of the first non-extinct type 1 cell, ℙ (*σ*_1_ *> t* | Ω_∞_), where *σ*_1_ represents the birth time and Ω_∞_ represents the event that the population does not go extinct, can be characterized as a function of the rate of beneficial gains *u*_1_ and growth parameters of type 1 and type 0 cells, respectively (Lemma 1). 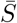 is thus,

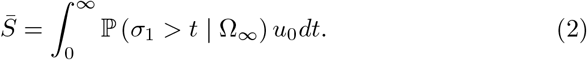

We explored various choices of growth parameters that capture the fitness difference between type 0 (with GD) and type 1 cells (with advantageous post-GD gain). Notably, lowering the fitness level of type 0 cells delays the birth of the type 1 cell (Figure 4C), conditioned on a fixed net growth rate of the type 1 cell. Accordingly, the post-GD gains become abundant with a bell-shaped distribution when the fitness difference is large. These model predictions are consistent with timing patterns in tumors with early GD (Figure 4A). Therefore, early GD observed in many patient tumors suggests that late-appearing gains may confer additional advantages for promoting the expansion. We note that the prolonged period of post-GD evolution could also be attributable to a lower rate of beneficial post-GD gains (Figure 4C). Nevertheless, the beneficial gain inevitably appears to be a late event since passenger gains introduced after the beneficial one most likely will be undetectable. The inability of passenger variants to become dominant themselves ensures such an outcome. Therefore, a long waiting time after GD suggests that the late beneficial gain becomes a rate-limiting event.

### 2.4 Ways evolving CN gains contribute to fitness increase and mutational diversification

As SCNAs have a global impact on gene expression in cancer [42], the evolving CN gains potentially affect dosage-sensitive genes whose gains have a functional impact. In the OS and BRCA tumors, as the CN evolves, we can see an enrichment of putative dosage-sensitive genes that are in pathogenic CNV peak regions derived from dbVar [43, 44] (Figure 5A). Notably, we observed a similar enrichment for genes involved in sustaining proliferative signaling: one of the most fundamental capabilities of cancer cells [1]. *MYC, EGFR* and *KIT* are among such genes with late gains in both OS and BRCA, emphasizing their ability in stimulating cell multiplication in multiple tumor types.

**Fig. 5.**
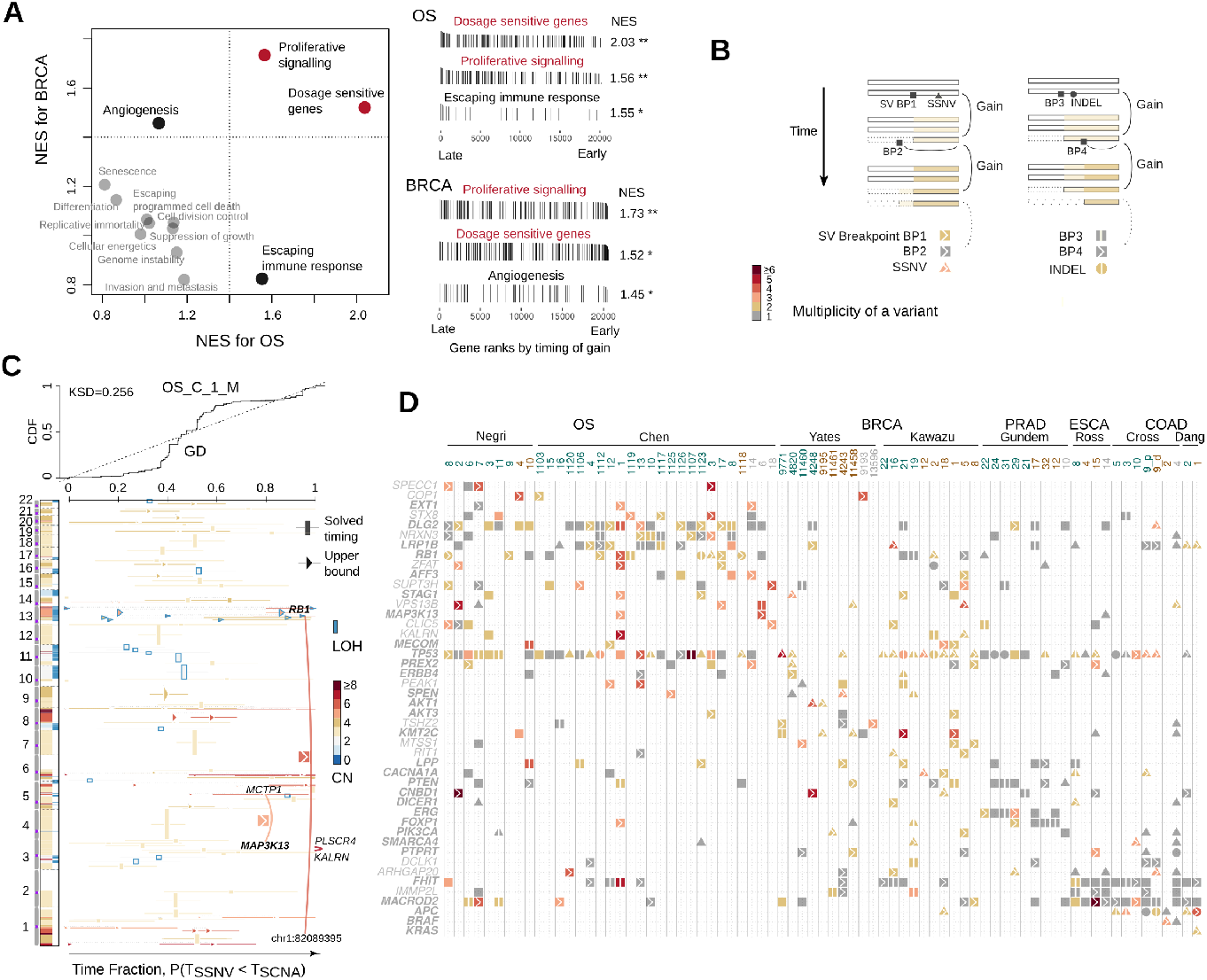
Ways the late CN gains contribute to the fitness of cancer progenitor cell. (A) The gene set enrichment analysis (GSEA) was performed on the gene list ranked by the averaged CN arrival time for BRCA and OS tumors, respectively. The scatter plot on the left shows the normalized enrichment scores (NES) for each set of cancer census genes belonging to the predefined cancer hallmarks by COSMIC database. The vertical bars on the right panels visualize the timing-ranks of genes that belong to the highlighted gene sets. (B) A cartoon illustrates the multiplicity increase of an early sequence variant due to the inclusion of that variant by a late CN gain, with annotations indicating the type of variants (symbol shapes), level of multiplicity (color hues) and the variants’ association with a late gain (right arrow) or an early gain (vertical bar). (C) The SCNA timing plot of an example OS tumor similarly arranged as in Figure 3, with additional links and symbols highlighting the SV breakpoints in known cancer genes that are amplified by late gains. (D) The matrix plot demonstrates genes with recurrent somatic variants and their multiplicity across the five tumor types. Names for known cancer genes are in bold. Genes with variants showing higher multiplicity levels than gene TTN are also included. Symbol annotations are the same as in (B).

The evolving gains could amplify the impact of early functional variants by increasing their multiplicity (Figure 5B). Such a mechanism potentially affects SV breakpoints in known oncogenes (e.g., *MAP3K13, MECOM* and *PREX2*), breakpoints in genes known to be involved in oncogenic fusions (e.g., *AFF3, LPP* and *ERG*), and simple mutations in oncogenes (e.g., SSNVs in *SMARCA4* and *CACNA1A*), see Figure 5C, D. *MAP3K13* had been shown to promote tumor growth in high *MYC* -expressing cells [45, 46], a similar context as in the OS [37]. We note that highly mutated tumor suppressor genes (TSG), such as *TP53, RB1* and *APC*, also have their early mutants duplicated or amplified (Figure 5D). Whereas these are presumably inactivation variants, the retaining of multiple copies of the variants could suggest different roles that remain unclear, such as a potential gain-of-function of APC mutants in COAD [47]. The fact that SRE requires the duplication of these early variants, rather than starting immediately upon acquiring a single copy of these variants, suggests that late-appearing gains could cooperate with the early variants to promote tumor expansion. On the other hand, late SV breakpoints (at single copy state) coupling evolving gains are prominent in genes located in common fragile sites, e.g., *FHIT* and *MACROD2*. Late alterations of these genome “caretakers” could facilitate further genome evolution and expedite clonal expansion [48, 49].

Lastly, the quantitative relation between SCNA evolution and SSNV accumulation, the rationale behind our timing method, implies that SCNA gains bolster mutational diversification between sub-populations during tumor growth. In principle, the higher the truncal CN state of a genomic segment, the higher the mutational divergence between subclones for the corresponding locus. As tumor expands, genomic regions at distinct SCNA states would accumulate SSNVs at different rates, leading to the heterogeneity of the SSNV burden along the genome. For example, when comparing two samples of a tumor, the sample-specific SSNVs are more abundant for regions with higher CN states (Figure 6A, B). Notably, the overall CN state affects the structure of phylogenetic trees, i.e., it explains more than 50% of the variance of the relative branching distance measured by SSNVs in COAD and PRAD patients, where extensive multi-region sampling is available (Figure 6C). Furthermore, continued evolution of SCNAs between subpopulations would also alter the SSNV divergence. For example, the SSNVs divergence is particularly enlarged for regions showing different CN states between the two samples (Figure 6B). As such, increased SSNV diversity in regions with CN gains provides more materials for further selection within the expanding cell populations.

**Fig. 6.**
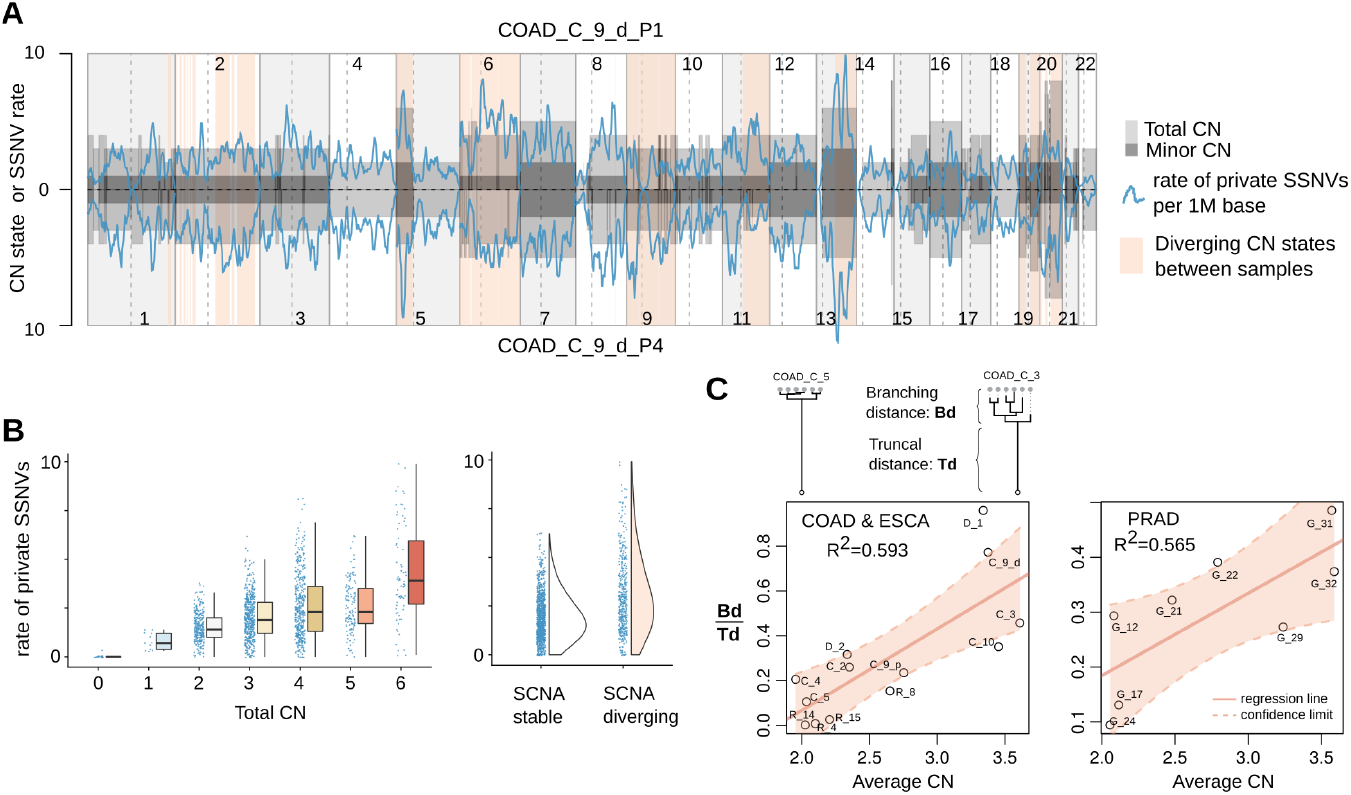
The effect of SCNA on SSNV diversification during tumor expansion. (A) The rate of sample-private SSNVs when comparing two samples of a COAD patient tumor. The segmental CN states (total and minor CN) along the autosomal chromosomes for the two tumor samples are shown as gray rectangles above and below the x axis. The rate of sample private SSNVs (per million base pairs, blue line) fluctuates with the CN states. Genomic regions with different CN states between the two samples are in light red background. (B) The box plots on the left panel illustrate the rate of private SSNVs in sample P1 detected in regions at a given total CN state. The half-violin plots on the right panel demonstrate such a rate for regions showing stable or diverging CN states between the two samples. (C) The branching distance relative to truncal distance in a tumor’s phylogenetic tree was calculated for each of the COAD, ESCA and PRAD tumors to evaluate the correlation with the averaged CN of the corresponding tumor samples. Annotations show the percentage of variance explained by a linear regression model.

## 3 Discussion

The major impediment to devising better therapies for tumors with the CIN phenotype is its highly complex somatic genome. In this study, we have created a computational framework for measuring the arrival and initiation time of SCNAs during the somatic evolution of the MRCA of tumor sample(s). By applying this method on WGS data of patient tumors, we have found that late truncal CN gains close to the most recent clonal expansion are common across multiple tumor types. Mathematical modeling predicts that these late evolving gains could contain rate-limiting driver events, promoting the tumor growth. As CN gains increase the gene dosage and early functional variants, we further demonstrated that an integrated analysis of SCNA timing, SV breakpoints and simple mutations has a strong potential for isolating the functional effect of specific genomic aberrations.

Early genomic changes are presumably beneficial for tumor initiation[27], but it is unclear the effect of late truncal events. Here we have provided evidence that gains occurring later in the somatic evolution, i.e., close to MRCA, can also be beneficial. The simplified two-event cancer development model posits that the cancer-initiating event is followed by the promoting event [50]. We reason that the evolving CN gains might render the progenitor cell capable of “self-promoting,” as they act similarly as a tumor “promoter” by (a) increasing the dosage of genes causing sustained proliferative signaling; (b) amplifying the mutant allele with early initiating driver variants; and (c) accelerating the accumulation of further genomic alterations. As both the early and late CN alterations could confer fitness advantages, chromosomal regions with SCNAs initiated early and arrived late, i.e., showing repetitive gains accompanying the entire course of the somatic evolution, could function as copy number “addictions.”

GD, a landmark event in CN evolution, has context-dependent fitness effect. The punctuated CN gains successfully induced the SRE in tumors that underwent a late GD. By contrast, for many other tumors, especially osteosarcoma, GD was followed by additional CN gains that produces the MRCA. GD could tolerate the occurrence of deleterious passengers [51]. However, simply escaping purifying selection was not sufficient to drive the ultimate outgrowth, at least not in the tumors with post-GD gains, where some chromosomal regions can reach higher CN states. Alternatively, GD may create an inflated genome space, accelerating the accumulation of driver alterations. As GD itself affects many genes, regions with pre- and/or post-GD gains could serve as a reduced search space for CN drivers.

Our method is applicable to a wide range of SCNAs, yet it is still challenging to analyze extremely high CN states (i.e. above eight). We note that regions with such a high CN likely evolve over time, such as the unequal segregation of extra-chromosomal oncogene amplifications [52, 53]. As such, late-arriving changes are expected for these amplified regions. Some focal high-level gains could involve small segments where the number of SSNVs is inadequate for calculation. This problem can be mitigated by borrowing information from nearby segments with the identical CN state. This strategy is applicable to synchronized SCNAs, such as chromothripsis [54, 55]. In addition, our analysis may have missed some late-appearing SCNAs due to overestimation. This is particularly evident for bi-allelic gains (4:2), where the upper bounds overestimate the actual timing (Supplementary Figure 1). Furthermore, deletion was not modelled as it is unidentifiable [18]; by comparing CN profiles between subpopulations, however, it is possible to study deletions during tumor expansion.

Our findings also illustrate the existence of a fundamental connection between CN evolution and SSNV diversity, which can explain the positive correlation between aneuploidy and mutational burden when excluding hyper-mutated tumors [56, 57]. Such a connection also indicates that we need to account for the dynamic nature of ongoing SCNAs when measuring subclonal evolution, which remains a challenge [58]. Finally, our results suggest that much can be gained by including the SCNA arrival time in studying tumor evolution, thereby shifting focus on exclusively early drivers to the evolving genomic events that affect the rate of tumor progression.

## 4 Methods

### 4.1 Somatic variant calling from WGS data

Raw WGS data in bam or fastq formats were downloaded from public databases provided by the original publications (Table 1). The cumulative read depth distribution along the human genome (hg38) and the tumor purity and ploidy for each sample are illustrated in Supplementary Figure 7, 8, and 9. We have extended our existing pipeline, which had achieved a balance in sensitivity and specificity in detecting SSNVs by borrowing information across multiple samples[60], to allow the detection of clonal SCNAs and the breakpoints of structural variants.

#### SSNVs and INDELs

Analysis-ready read alignment bam files (against hg38) were generated according to the best practices, including indel realignment, base recalibration and flagging of duplicated reads. Raw candidate variants were produced by MuTect (v1.1.7) [3]. To reduce the false-positive rate due to misalignments or other technical artifacts and to salvage the variants that may be missed due to uneven read coverage between samples, the alignment features surrounding each candidate variant were collected for each sample. The heuristic-based criterion for the read alignment patterns was adopted to refine and variant calls as detailed previously [60]. Small insertions and deletions were called by using Strelka (v1.0.15)[61].

#### SCNAs

Copy number and tumor purity were estimated by using TitanCNA (v1.26.0)[25]. Germline heterozygous SNVs used as input to TitanCNA were identified using Samtools (v1.5)[62] and subject to the same filtering strategy as was applied to SSNVs. The one-clone solution reported by TitanCNA (i.e., the sample is dominated by a clone with an SCNA profile along the genome) globally fit the data of the read coverage and allelic imbalance well, with a few exceptions for which the two-clones solution are necessary to explain the data of specific genomic regions.

#### SVs

We incorporated two distinct SV calling tools relying on orthogonal approaches, i.e., DELLY (v0.7.8, abnormal read pair and split-read analysis)[63] and GRIDSS (v2.10.1, local assembly based algorithm)[64]. We focused on the SV breakpoints found by both tools, as these shared calls generally have higher quality (e.g. with higher breakpoint confidence) than those unique to each tool (Supplementary Figure 10). SV breakpoints were annotated with AnnotSV[65].

### 4.2 Analysis of genomic divergence

#### SCNA divergence

When multi-samples are available for a patient, the truncal and private SCNAs were identified as follows: (1) we partitioned the genome into disjoint segments by considering all the SCNAs called from the samples of the patient; (2) for each segment, we calculated a generalized likelihood ratio statistics for the comparison between two samples. The statistics is the ratio of the values of the likelihood function (the probability of observing the read depth ratio and B-allele frequency for SNPs in the region) evaluated at the maximum likelihood estimation in the sub-model (two samples have the same CN profile) and at the maximum likelihood estimation in the full-model; and (3) the statistics converges weakly to a random variable with chi-square distribution and thus can be used to determine if a segment shows significantly different SCNA states between the two samples. The term “truncal SCNAs” refers to SCNAs that exhibit no difference in pairwise comparisons.

#### Sample phylogeny

We applied Treeomics (v1.7.13) [66] to construct sample phylogenies from SSNV data. Treeomics takes into account the uncertainty due to purity differences and variations of read depth on the SSNV loci to derive robust sample phylogenies.

#### Clonality, multiplicity of SSNVs and SV breakpoints

SSNVs were classified as public (present in all tumor cells) or private according to the criterion as previously described for multi-sampling data [60]. In individual samples, clonal SSNVs were identified as those with the 95% confidence interval of cancer cell fraction (CCF) covering 1 [60]. We focused on the public SSNVs (multi-sampling) and clonal SSNVs (single sampling) for the timing analysis. For SSNVs or SV breakpoints existing in an SCNA region, we applied a binomial model to calculate the maximum likelihood estimates of the number of segment copies containing that variant as previously described [21].

### 4.3 Inferring the arrival and initiation time of SCNAs

#### 4.3.1 Allele state distribution of SSNVs for a clonal SCNA

For SSNV *i* in an SCNA region (with CN configuration of *N*_*t*_ : *N*_*b*_ and *M* ≥ 10 SSNVs in total), we obtained from WGS the read counts carrying the mutant allele *m*_*i*_ out of the total number of reads *d*_*i*_. Expectation Maximization algorithm was used to estimate the proportion of SSNVs at each possible allele fraction, i.e. a vector *q* that gives the probability of randomly acquired SSNVs in this region having an purity-adjusted allele frequency (*f*_*i*_ = *a*_*j*_) for each possible allele state 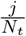. The rationale behind this is that the likelihood function of observing a particular SSNV data is related to the probability vector *q* as well as tumor purity. The log-likelihood is given by,

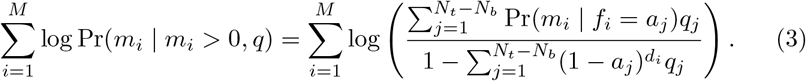

#### 4.3.2 Estimating the upper bounds of initiation and arrival time

For a genomic region affected by SCNA gains, let A be a possible history matrix with entry *A*_*jk*_ representing the number of segment copies in stage *k* that result in a final allele state *a*_*j*_. Let *q* be a vector with entry *q*_*j*_ representing the probability of a randomly acquired SSNV in this region having allele frequency *a*_*j*_. For single and double gains (*Nt* : *Nb* at 3:1, 2:0, 4:1 or 3:0), the initiation time (*t*_0_) and arrival time (*t*_*K*_) are directly solved because matrix A is unique and invertible. For other SCNAs, `Butte` uses linear programming to obtain the upper bounds of timings across all possible history matrices for the corresponding CN configuration (Supplementary Figure 11). Let *s* denote the vector of the column sum of matrix *A*. Let *t* denote the relative time vector in which every component is a time fraction of the corresponding copy number state. We solve the following optimization problem by linear programming:

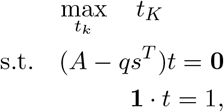

where *t*_*K*_ is the last element in vector *t*. The maximum value of *t*_*K*_ gives us an upper bound of the arrival time given *A*. For upper bounds of initiation, we instead maximize *t*_0_ which is the first element in *t*. To tolerate noise in the allele state distribution estimated from sequencing data, we add a slack variable on each capacity constraint, having a penalty cost of 100. The confidence intervals of the estimated upper bounds were calculated through bootstrapping the SSNV data.

### 4.4 Mathematical modeling of the late evolving gains

Consider two contrasting models based on multi-type branching processes with mutations. In both models, the tumor grows from a single tumor-initiating cell which just acquired GD. During the tumor’s progression, cancer cells accumulate mutations (post-GD gains).

In the first model, all mutations are passenger mutations. Therefore, all cancer cells give birth at a rate of *a*_0_ and die at a rate of *b*_0_. The net growth rate is *λ*_0_ = *a*_0_ − *b*_0_ *>* 0. Neutral mutations occur at rate *u*_0_ per unit time throughout the lifetime of a cell, and each mutation is distinct (the infinite-sites model of Kimura [67]). We can obtain that the number of mutations accumulated before the first cell division, which results in two descendants with infinite lineage, follows a geometric distribution with parameter 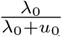 and mean 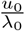 (conditioned on the non-extinction of the cancer cell population). We then investigated the number of mutations which are shared by more than 90% of the total population (we refer to them as dominant mutations). Gunnarsson and his co-authors [41] derived exact expressions for the expected SFS of a cell population that evolves according to a branching process. We utilized their results on the skeleton subpopulation (see Appendix C of [41]) — cells with an infinite line of descents which determines the high frequency spectrum — to express the expected number of dominant mutations 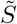 when the tumor reaches a fixed size *N* as

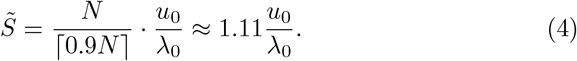

In the second model, the tumor-initiating cell and its descendants with only passenger mutations form the type 0 population. Type 0 cells give birth at a rate of *a*_0_ and die at a rate of *b*_0_. The net growth rate is *λ*_0_ = *a*_0_ − *b*_0_ *>* 0. Type 0 cells mutate to type 1 cells at a rate of *u*_1_. Type 1 cells give birth at a rate of *a*_1_ and die at a rate of *b*_1_. The net growth rate is *λ*_1_ = *a*_1_ − *b*_1_ *> λ*_0_. Both type 0 and type 1 cells accumulate passenger mutations at a rate of *u*_0_. Due to the selective advantage type 1 cells have over type 0 cells, a tumor at detection is most likely to be dominated by type 1 cells. As a result, it suffices to focus on mutations shared by the majority of type 1 cells. For tractability, we investigated the number of passenger mutations accrued in the first type 1 cell with infinite lineage. In Lemma 1, we obtained the distribution of the time to the first such type 1 cell conditioned on the non-extinction of the tumor.

#### Lemma 1

*Let σ*_1_ *denote the time of occurrence of the first type* 1 *cell that gives rise to a family which does not die out, and let* Ω_∞_ *denote the event of non-extinction of the tumor. Then*

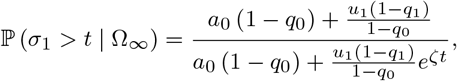

*where*

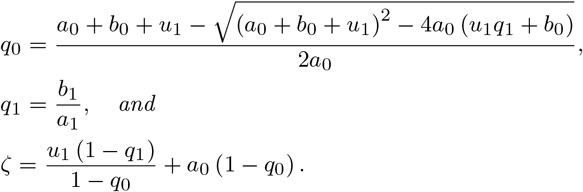

With Lemma 1, we can obtain the expected number of passenger mutations accumulated in the first type 1 cell with infinite lineage, denoted by 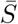:

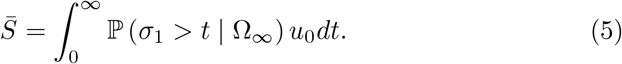

With (4) and (5), we obtained that the expected number of dominant post-GD gains in the subpopulation generated from the first type 1 cell with infinite lineage is 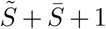, where the last 1 represents the number of post-GD driver mutation. Proof for Lemma 1 and details of (4) can be found in Supplementary Methods.

## Acknowledgments

This study uses the computing resources in Minnesota Supercomputing Institute. This manuscript was prepared using limited access datasets obtained from BC CANCER (OS N), Queen Mary University of London (COAD C), the Cancer Genome Project in Wellcome Sanger Institute (BRCA Y), and does not necessarily reflect the opinions or views of the corresponding provider institutions. The OS C dataset was generated by the St. Jude Children’s Research Hospital – Washington University Pediatric Cancer Genome Project. The BRCA K dataset was originally generated by research led by Dr. Masahito Kawazu and available at the website of the National Bioscience Database Center (NBDC; http://biosciencedbc.jp/en/) of the Japan Science and Technology Agency (JST). We thank International Cancer Genome Consortium (ICGC) for providing the access to the PRAD G dataset (originally produced by Cancer Research UK Prostate Cancer Group Study) and ESCA R dataset (by the Oesophageal Cancer Clinical and Molecular Stratification Study Group). We would also like to acknowledge the database of Genotypes and Phenotypes (dbGaP) and Dr. Ryan C. Fields for producing the COAD D dataset (phs001722.v1.p1) which was supported by Siteman Cancer Center Investment Program. We thank Dr. Scott Dehm and Boyang Liu for commenting on the manuscript. We thank Einar Gunnarsson for sharing his proof ideas on the skeleton of the branching process.

## Code availability

Code for `Butte` (computational framework to estimate SCNA arrival and initiating time from WGS data) and the mathematical modeling is available at https://github.com/SunPathLab/Butte/

## Author contributions

R.S. and Z.W. designed the study. R.S., Y.X., and Z.W. developed the algorithms. Z.W., R.S., and A.N.N. constructed mathematical models and performed simulation studies. R.S., Y.X., Z.W. and L.M. performed the analysis of WGS data and visualized the results. R.S., Z.W., Y.X., N.M. and J.M.S interpreted the results and wrote the manuscript. All authors reviewed and provided feedback on the manuscript.

